# Continuous Capture of recombinant AAV Particles Using Twin-Column CaptureSMB

**DOI:** 10.64898/2026.06.12.731701

**Authors:** Julia M. Müller, Daniela Tobler, Jules Bühler, Damian Hauri, Richard Plieninger, Sven Göbel, Efe Saygili, Yuki Higuchi, Ryosuke Takahashi, Sebastian Vogg, Thomas Müller-Späth, Thomas K. Villiger

## Abstract

Recombinant adeno-associated viruses (rAAVs) have gained increasing importance in gene therapy due to their safe and precise gene delivery. However, certain indications require substantially higher vector doses, pushing manufacturing capacity and cost of goods (COG) to its limits. In this study, we present for the first time a continuous twin-column capture process (CaptureSMB) enabling direct purification of rAAV5 from unprocessed perfusion harvest without prior concentration or processing. This approach differs fundamentally from conventional batch workflows which typically mandate clarification and concentration before affinity capture and offers a novel process integration in viral vector manufacturing.

A single-column batch capture process was developed first and subsequently compared to continuous CaptureSMB configurations. Optimized CaptureSMB operation achieved consistent yields over four cycles, with recoveries exceeding batch operation (+ 14.3%) with concomitant higher productivity (+ 11.4%) and reduced buffer consumption (- 79.2%). Critical quality attribute analysis showed lower host cell protein levels and lower residual DNA in early CaptureSMB cycles, while full capsid ratios, thermal stability and transduction efficiency of rAAV5 particles remained unaltered across cycles and process modes.

These findings highlight that continuous twin-column CaptureSMB directly from perfusion harvest can not only improve yield and manufacturing efficiency but also maintain and in some respects enhance product quality. This novel strategy provides a promising route to address manufacturing capacity and cost challenges in rAAV gene therapy production.

## Introduction

The clinical application of viral vectors is expanding the biopharmaceutical repertoire for treating refractory genetic, oncological, and autoimmune diseases (Currier et al., 2025; George et al., 2017; Kuwahara et al., 2026; Mease et al., 2010; Mendell et al., 2017). Among these vectors, recombinant adeno-associated viruses (rAAVs) stand-out due to their favorable safety profile, highly specific delivery, broad tropism, and their therapeutic efficacy (Fischer et al., 2019; Han et al., 2024; Ling et al., 2023; Liu et al., 2025; Sobh et al., 2023).

Despite their promise, rAAV-based therapies face significant dose-challenges in certain indications. Local administration, as typically seen in ophthalmology, target restricted therapeutic areas and can be treated with relatively small vector doses. In contrast, systemic indications such as clotting disorders or Duchenne muscular dystrophy require widespread distribution throughout the body. The large quantities of rAAV viral genomes (vg) required in these cases place considerable pressure on manufacturing processes, which are still not fully established in the industry to the level of generic platform processes.

A conventional large-scale rAAV process typically begins with transient triple transfection in a batch bioreactor (Chahal et al., 2014; Grieger et al., 2016). This is followed by cell lysis, nuclease digestion, and depth filtration. These steps are associated with considerable costs, primarily due to the high expenses associated with the large quantities of plasmid and nuclease needed for efficient host cell DNA digestion (Lyle et al., 2024; Thakur et al., 2025a). In batch processing, media is consumed in a single production run, with costs largely determined by the size of the bioreactor and the cell density achieved. Following lysis and digestion, tangential flow filtration (TFF) is commonly used as concentration step (Lock et al., 2010; Turco et al., 2023; Žigon et al., 2022). This step reduces the loading time for the first chromatography step and allows for buffer exchange to remove impurities, such as digested nucleic acid fragments and host cell proteins (Cardoso et al., 2026; Chaubal et al., 2026; McCarney et al., 2025; Miyaoka et al., 2023). However, implementing TFF prior to affinity capture introduces additional complexity: it requires extra equipment, increases the manufacturing footprint, and potentially compromises product yield or quality due to shear forces (Leach et al., 2025; McCarney et al., 2025; Miyaoka et al., 2023; Picciano et al., 2025). Sequential lysis, nuclease digestion, depth filtration, and ultrafiltration in standard rAAV processing trains can result in losses of 20 - 80% of upstream vector mass before capture and pose risks such as irreversible filter cake fouling and shear-induced aggregation (Chaubal et al., 2026; Leach et al., 2025; Picciano et al., 2025).

After affinity chromatography, anion-exchange (AEX) chromatography is typically used to improve purity and enrich the proportion of full rAAV particles in the final product (Duzenli and Aslanidi, 2025; Kurth et al., 2024; Lavoie et al., 2023; Thakur et al., 2025b). Although these successive steps are effective individually, overall process yields and purity are often suboptimal, with yields reaching as low as 20 - 35% (Florea et al., 2023; Hebben, 2018; Rieser et al., 2021; Tomono et al., 2018; Wright et al., 2005). For batch processes, media costs are generally lower than in perfusion, but reagents and equipment still represent significant cost drivers. Despite notable progress in product development and clinical success, the long-term viability of rAAV therapeutics depends on optimizing manufacturing strategies to lower costs and improve accessibility of rAAV based therapies (Emily et al., 2019; Sarkis et al., 2023; Thakur et al., 2024).

Substantial technical innovations aimed at improving efficiency and productivity have emerged (Chen et al., 2024; Mendes et al., 2022b; Park et al., 2024; Ren et al., 2025; Thakur et al., 2025a). One promising approach is the application of continuous chromatography. This technique offers several advantages over conventional batch chromatography, including increased productivity, reduced buffer consumption, and improved yield while maintaining a high purity in polishing stages (Angarita et al., 2015; Müller-Späth et al., 2010). Recent studies have begun to explore the implementation of continuous capture chromatography in rAAV manufacturing (Mendes et al., 2022a; Müller et al., 2025; Neto et al., 2025; Ossi et al., 2025), suggesting its potential to enhance existing purification strategies.

Previous studies on continuous capture have used multicolumn approaches using three- or four-column configurations in combination with pre-purified rAAV feed, involving DNA digestion and TFF concentration. Most established studies have relied on the AAVX affinity resin (Florea et al., 2023; Mendes et al., 2022a; Neto et al., 2025), which effectively binds multiple rAAV serotypes, yet has notable drawbacks in crude feed capture. Reports indicate fouling due to host cell protein, nucleic acids and lipids, leading to fast clogging and reduced chromatography resin lifetime (Florea et al., 2023; Leibiger et al., 2024; Soni et al., 2025, 2024). As a result, binding performance in capture chromatography can significantly decrease when crude harvest is loaded without pre-treatment, such as DNase digest, and enzymatic DNA removal (Florea et al., 2023; Mendes et al., 2022a). Consequently, the short resin lifespan and the requirement for feed pretreatment make the process less efficient and more complex to operate.

In this study, we demonstrate a continuous twin-column CaptureSMB process for direct capture of rAAV particles from untreated harvest of a perfusion bioreactor. To the best of our knowledge, no reports have described the application of a twin-column counter-current CaptureSMB process for the direct capture of viral vectors. Unlike previous studies, our approach bypasses DNA digestion, depth filtration, TFF upconcentration and buffer exchange, thereby eliminating a major source of yield loss and process complexity identified in conventional rAAV capture workflows. Our results show that this process achieves high recovery and maintains product integrity while substantially reducing buffer consumption, providing a key enabling step toward fully integrated continuous manufacturing of viral vectors.

## Material and Methods

### rAAV Production

HEK293F cells (Viral Production cells 2.0, Thermo Fisher, Waltham, Massachusetts, USA) were cultivated in HyClone Peak Expression Medium (Cytiva, Marlborough, Massachusetts, USA) and expanded in shake flasks in an incubator (Multitron, Infors HT, Bottmingen, Switzerland) at 36.5 °C at 5% CO_2_ and 120 rpm (50 mm orbital throw). Once the desired cell concentration was reached, a transient triple transfection was performed. All three plasmids used pHelper (pAdDeltaF6, 112867), pRepCap (pAAV2/5, 104964), and pTransfer (pAAV.CMV.PI.EGFP.WPRE.bGH, 105530) were obtained from Addgene and amplified and purified in-house. The plasmid DNA was combined at a defined molar ratio of 1.2:2.8:1 (pHelper: pRepCap: pTransfer, 2 μg of total plasmid DNA per million cells), and complexed with polyethylenimine Max (PEI Max, Polysciences) in a 2:1 ratio.

Following transfection, the cells were seeded into a Labfors 5 bioreactor (Infors HT, Bottmingen, Switzerland) at a viable cell concentration (VCC) of 1.5 × 10^6^ cells/mL and a working volume of 1.6 L. The bioreactors were maintained at 36.5 °C with dissolved oxygen controlled at 50% and stirring at 270 rpm. Perfusion was initiated at a VCC of 3 × 10^6^ cells/mL using a 0.19 m² hollow-fiber module (0.4 µm, Asahi Kasei, Tokyo, Japan) operated in reverse tangential flow mode (Romann et al., 2024). To obtain homogeneous feed, the perfused harvest of two perfusion runs was pooled and aliquoted. For the downstream experiments, the required material was thawed overnight at room temperature and 0.2 µm filtered into a sterile bottle. The feed bottle was connected to the chromatography system via a 0.2 µm filter (Supor KA02EAVP8G, Cytiva, Marlborough, Massachusetts, USA).

### Chromatography Equipment

The continuous twin-column chromatography system Contichrom® CUBE 30 (ChromaCon AG, Zurich, Switzerland) with the ChromIQ® version 9.0 operating software was used to perform the preparative chromatography runs. The conductivity and UV signal were monitored online after each column. UV detectors with a path length of 1.0 mm were used and the absorbance was measured at 260, 280, and 320 nm. Additionally, the pH was monitored online at the product outlet line.

### Batch Affinity Chromatography Method

For the capture of rAAV5, an AVIPure AAV5 affinity column (5 × 25 mm, 0.5 mL, Repligen 23051305), with an alkali-tolerant recombinant protein ligand coupled to a 50 µm agarose base matrix, was used. After equilibration with 10 column volumes (CV) of equilibration buffer (50 mM Tris-HCl (Sigma, 10708976001), 250 mM NaCl, 0.001% (w/v) poloxamer 188 (Thermo Fisher Scientific, J66087.36), pH 8.0), the perfusion harvest containing the secreted rAAV5 from was loaded onto the column with a residence time of 0.45 min (336 cm/h). After the load, a post-load wash (PLW) was performed for 10 CV using equilibration buffer. A low pH elution buffer (100 mM glycine (Carl Roth, 3187), 240 mM arginine (Carl Roth, 3144), 150 mM NaCl, 0.01% (w/v) poloxamer 188, pH 2.5) was used to elute the rAAV5 capsids, and the eluate was directly neutralized with 12% v/v neutralization buffer (1 M Tris-HCl, pH 10). The column was subsequently regenerated with 10 CV strip solution (1 M NaCl), 25 CV 0.1 M NaOH (Carl Roth, 9356) and another 5 CVs of 1 M NaCl before re-equilibrating the column with 10 CVs of equilibration buffer. Equilibration was performed at 250 cm/h, PLW at 100 cm/h, strip at 200 cm/h, cleaning-in-place (CIP) at 150 cm/h, and the elution at 75 cm/h. The column was loaded to its DBC at 1.5 or 1% breakthrough. These conditions served as reference for the CaptureSMB run.

### CaptureSMB

For the CaptureSMB run, two AVIPure AAV5 columns (5 × 25 mm, 0.5 mL, 50 µm, Repligen, 23051305) were used. CaptureSMB is a continuous chromatography process that is operated in a cyclic manner, where the columns switch between interconnected and batch set-ups by valve switching (**Figure 1**). To enter this cyclic phase, a start-up method is needed first to pre-load the first column in an interconnected state before the run enters the cyclic phase. During this phase, the column continues to be loaded in an interconnected state, where the second column captures rAAV particles that flow through the first column (**Figure 1A**). Once the first column is loaded to a DBC of 16% with perfused rAAV5 harvest (1.8 cp/mL), an interconnected PLW step is performed to flush the remaining load material present in the interstitial volume of the first column onto the second column. Afterwards, the columns are set to a parallel operation during which the first column undergoes elution and regeneration (**Figure 1B**). In the meantime, the second column is continuously fed with harvest at a reduced flow rate (270.5 cm/h) not to exceed the DBC of 1% breakthrough, which was defined for the loading in batch mode. Once the first column is fully regenerated and re-equilibrated, the first switch is completed. At this stage, the columns switch relative position and are set to an interconnected state again, where the regenerated column is in the downstream position capturing the flowthrough of the first column (**Figure 1C**). If this upstream column is again fully loaded and eluted (**Figure 1B**), both columns go back to their original position, and a full CaptureSMB cycle is completed. In total 4 cycles were performed in this first proof-of-concept. To finalize the run, a shutdown procedure was applied to elute the last column and recover the remaining rAAV5 particles from the column.

**Figure 1:**
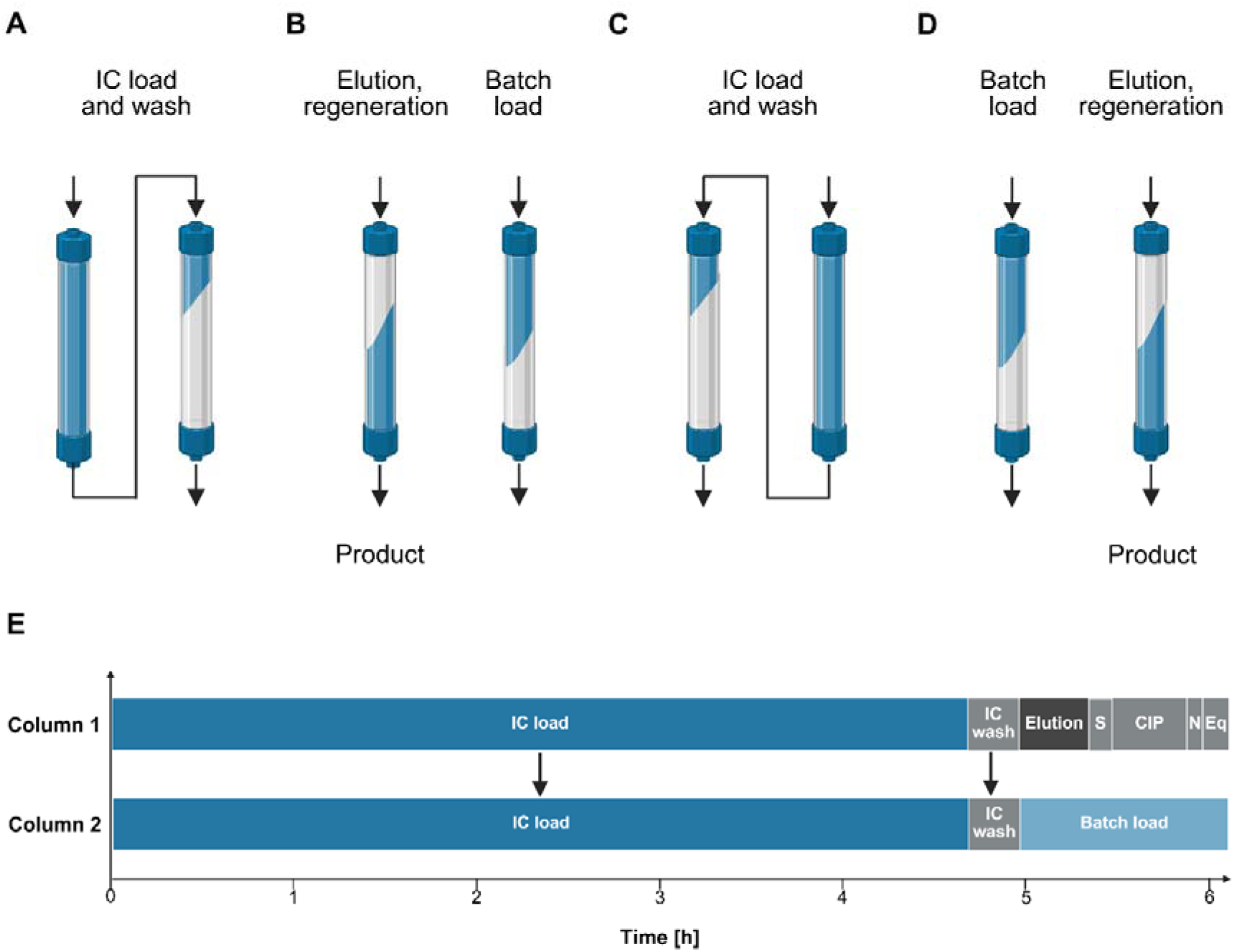
Schematic representation of the CaptureSMB. **(A)** In the cyclic CaptureSMB operation, the product is first loaded in an interconnected state (IC load), followed by an interconnected wash (IC wash). **(B)** After that, the valve positions switch the columns to a parallel operation, where the product is eluted from the first column followed by the column regeneration. The second column, in the meantime, is fed in batch mode (batch load). **(C)**, **(D)** As the first column is fully regenerated, the first switch is completed and the columns switch relative positions, starting the second switch of a cycle. **(E)** Time regime for a CaptureSMB switch. S: strip; CIP: cleaning in place; N: neutralization; Eq: equilibration. Created in BioRender. Müller, J. (2026) https://BioRender.com/gynqlj2

### Capsid Titer Measurement

rAAV capsid titers were quantitatively determined using the rAAV5 Xpress ELISA kit (PROGEN, PRAAV5XP) according to the manufacturer’s instructions. The different samples were diluted 1 × assay buffer to ensure that the capsid concentrations remained within the standard curve. The dilutions were optimized empirically and resulted in a 200- and 400-fold dilution for load samples, a 5- to 40-fold dilution for flow-through samples and an 8,000- and 16,000-fold dilution for eluate samples. Absorbance was recorded at 450 nm using a BioTek Epoch 2 microplate reader (Agilent Technologies).

### Anion Exchange High-Performance Liquid Chromatography (AEX-HPLC)

The full/empty ratio measurements were carried out on an Agilent 1260 Infinity II Bio-inert LC System (Agilent Technologies, USA) in combination with a BioPro IEX QF column (YMC CO., LTD., Japan, 30 × 4.6 mm, 5 µm, QF00S05-0346WP) maintained at 25 °C. The injection volume was set to 5 µL, and the flow rate to 0.6 mL/min. The maximum operating pressure was limited to 80 bars, with a total runtime of 15 min per analysis. The mobile phases consisted of buffer A (20 mM Tris, pH 8.5 ± 0.05) and buffer B (20 mM Tris, 120 mM MgCl₂, pH 8.5 ± 0.05). The separation was performed using a stepwise gradient starting at 95% buffer A and 5% buffer B, which was held for 1 min. The composition was then adjusted to 83% buffer A and 17% buffer B and maintained for 1 min, followed by a gradual increase to 35% buffer B over the next 6 min. Subsequently, the column was washed with 100% buffer B for 1 min before being re-equilibrated to the initial conditions (95% buffer A) for 2 min prior to the next injection. UV detection was performed using a DAD detector at wavelengths of 230, 260, 280, and 320 nm (attenuation: 1000 mAU; zero offset: 5%). Fluorescence detection was conducted at excitation/emission wavelengths of 280/331 nm with a peak width of 2.31 Hz (attenuation: 100 LU; zero offset: 5%).

### Bicinchoninic Acid (BCA) Protein Assay

The total protein content of process samples was determined using the Pierce BCA Protein Assay Kit (Thermo Scientific, 23227) according to the manufacturer’s instructions. A volume of 25 µL of each sample or standard were mixed with 200 µL working reagent and incubated at 30 min for 37 °C. The absorbance was measured at 562 nm using a BioTek Epoch 2 microplate reader without prior dilution. The total protein content was normalized to the full viral particles.

### Residual DNA Measurement

Residual dsDNA was quantitatively determined using the HyperLight FluorGreen dsDNA Assay kit (APExBIO, K1603) according to the manufacturer’s instructions. Optimal dilutions with dsDNA HS Buffer were determined empirically to ensure that the absorption signal was within the calibration curve. Load samples were diluted 100-fold and eluate samples 10-fold. The samples were then mixed with an equal volume of dye working solution before measuring the fluorescence intensity (Ex = 480 nm; Em = 520 nm) on a BioTek Synergy H1 multimode microplate reader (Agilent Technologies). The residual DNA content was normalized to the full viral particles.

### Transduction Efficiency Measurement

The functional activity of the eluted rAAV5 was determined using a GFP-based transduction assay. Adherent HEK293 cells (ATCC; CRL-1573) were seeded at 3 × 10^5^ cells/mL in 24-well plates and incubated for 24 h. Subsequently, the VCC was determined using 3 of the 24 wells. To ensure a consistent multiplicity of infection (MOI), eluate samples were first normalized to their full rAAV titer. The samples were then diluted in DMEM + 10% heat-inactivated FCS to a final volume of 150 µL and added to the cells at an MOI of 5 × 10^3^ vg/cell. Following a 4 h incubation, 500 µL of fresh DMEM medium +10% heat-inactivated FCS medium were added. The cells were incubated for another 72 h, before harvesting and straining (100 µm, Corning, 431752). For the analysis of GFP positive cells by flow cytometry (Sony SH800ZFP, San Jose, California), only wells with 2-20% GFP-positive cells were used. Consequently, the specific activity was calculated by normalizing the transducing units (TU/mL) by the amount of full AAV determined by AEX-HPLC and capsid measurement.

### Polydispersity and Thermal Stability Measurement

Polydispersity index (PDI) and thermal stability were measured with the Prometheus Panta (NanoTemper Technologies, Germany) using dynamic light scattering (DLS) by nano differential scanning fluorimetry. For the thermal stability measurement, the samples were heated from 20 to 99 °C at a constant rate of 1 °C/min. The melting temperature (T_m_) was determined from the inflection point of the intrinsic fluorescence ratio (350/330 nm).

### Process Performance Evaluation

Batch and CaptureSMB processes were evaluated using three key metrics, namely yield, productivity, and buffer consumption. Yield (*Y*) is defined as the ratio between the recovered product mass (*m^cp,recovered^* and the product mass supplied through the feed (*m_cp,feed_*) using the capsid measurements:

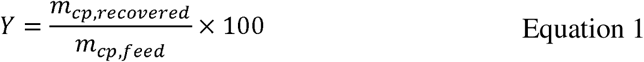

Productivity (*Pr*) is defined as the ratio of the entire product mass recovered *m^cp,recovered^* and the total process duration (*t*) multiplied with the total column volume (*mCV_Col_*):

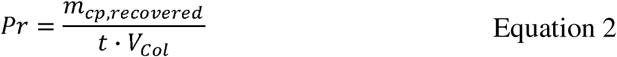

Buffer consumption (*BC*) is defined as the ratio of the total buffer used *V^buffer^* and the total recovered product mass:

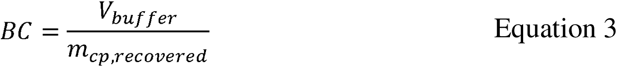

## Results

Recombinant AAV5 (rAAV5) was produced in HEK293F suspension cells via transient triple transfection and harvested using a 0.4 µm hollow-fiber membrane. Titers of rAAV in the upstream harvest varied between cultivations. To obtain a homogeneous feedstock, material from two perfusion cultivations was pooled, resulting in a final titer of 1.8 × 10¹¹ cp/mL. Additionally, a portion of harvest with the highest titer (4.8 × 10¹¹ cp/mL) was kept separate to enable breakthrough experiments at higher load levels. The pooled material was used for two breakthrough curves and the CaptureSMB run, while the high-titer material was used for two additional breakthrough curves. A batch capture process was developed using an AVIPureAAV5 affinity column and subsequently applied in breakthrough studies with both feedstocks at flow rates of 360 and 214 cm/h (**Figure 2**). The resulting empirical breakthrough data served as the basis for designing the continuous CaptureSMB process. Using the ChromIQ CaptureSMB wizard, the duration of batch load and integrated load phases was determined, yielding a utilized dynamic binding capacity (DBC) at 1.5% breakthrough for batch loading and at 17% breakthrough for interconnected loading (**Table 1**, design A).

**Figure 2:**
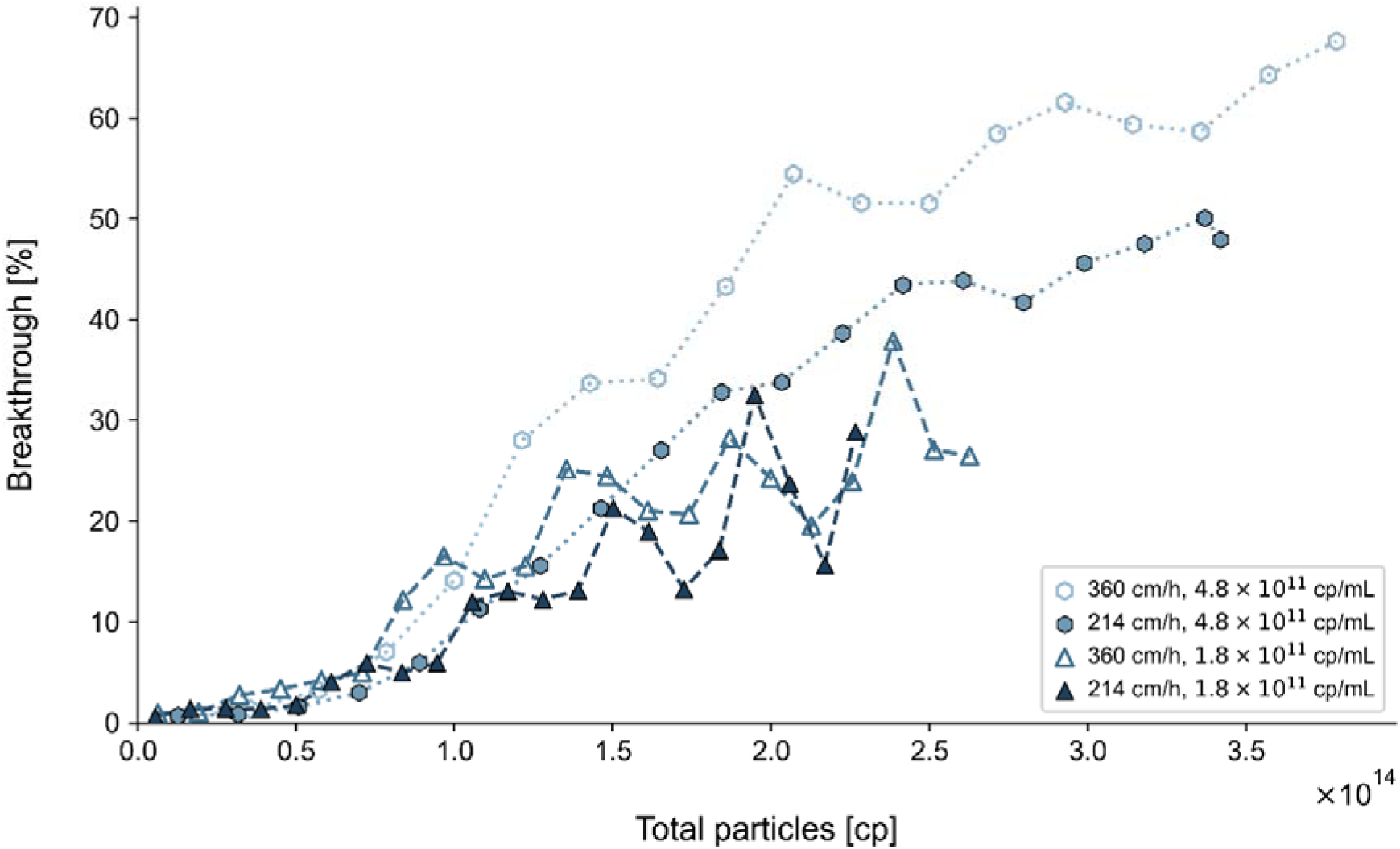
Breakthrough curves with percentage of rAAV5 breakthrough in the flowthrough for different feed concentrations and flow rates.

**Table 1:**
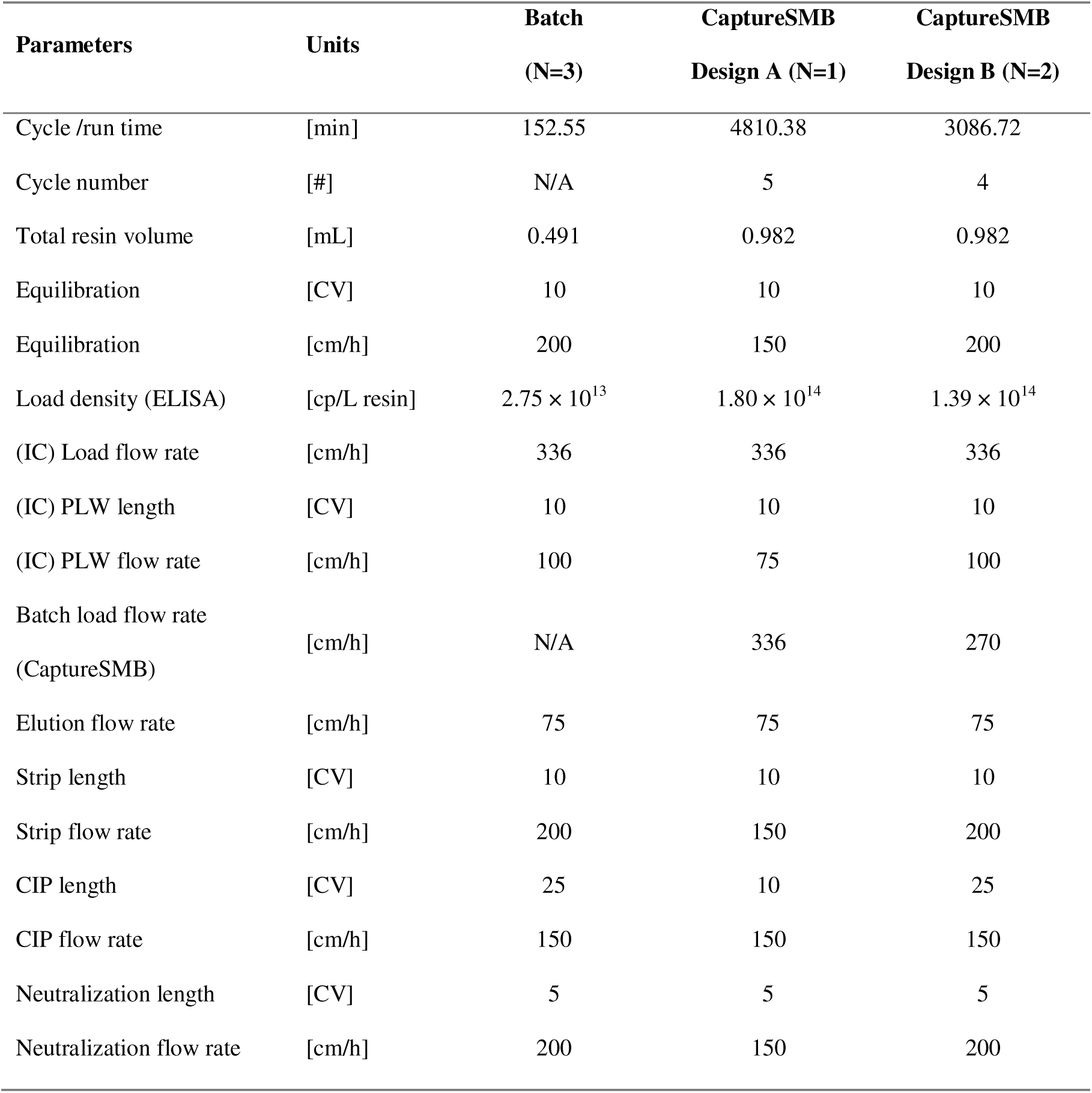
Process Parameters overview – Batch versus CaptureSMB. The overview of the process parameters is shown for conventional batch operation and the CaptureSMB designs A and B. IC: Interconnected.

| Parameters | Units | Batch | CaptureSMB | CaptureSMB |
| --- | --- | --- | --- | --- |
|  |  | (N=3) | Design A (N=1) | Design B (N=2) |
| Cycle /run time | [min] | 152.55 | 4810.38 | 3086.72 |
| Cycle number | [#] | N/A | 5 | 4 |
| Total resin volume | [mL] | 0.491 | 0.982 | 0.982 |
| Equilibration | [CV] | 10 | 10 | 10 |
| Equilibration | [cm/h] | 200 | 150 | 200 |
| Load density (ELISA) | [cp/L resin] | $2.75 \times 10^{13}$ | $1.80 \times 10^{14}$ | $1.39 \times 10^{14}$ |
| (IC) Load flow rate | [cm/h] | 336 | 336 | 336 |
| (IC) PLW length | [CV] | 10 | 10 | 10 |
| (IC) PLW flow rate | [cm/h] | 100 | 75 | 100 |
| Batch load flow rate<br>(CaptureSMB) | [cm/h] | N/A | 336 | 270 |
| Elution flow rate | [cm/h] | 75 | 75 | 75 |
| Strip length | [CV] | 10 | 10 | 10 |
| Strip flow rate | [cm/h] | 200 | 150 | 200 |
| CIP length | [CV] | 25 | 10 | 25 |
| CIP flow rate | [cm/h] | 150 | 150 | 150 |
| Neutralization length | [CV] | 5 | 5 | 5 |
| Neutralization flow rate | [cm/h] | 200 | 150 | 200 |

Evidence of column fouling was first observed through a gradual increase in column pressure from cycle to cycle (**Figure S1**) during the initial CaptureSMB process (design A). This pressure trend is further supported by declining yields and significant changes in peak shape (**Figure S2**), both of which suggest notable accumulation of foulants. These effects are likely driven by residual DNA and host-cell proteins in the feed material, which can impair column performance even in the absence of full cell lysis.

To mitigate fouling, a second configuration (design B) was implemented with lower resin utilization targets. Interconnected loading was reduced from 17% to 16% dynamic binding capacity (DBC), and batch loading from 1.5% to 1% (**Table 1**, design B). These adjustments shortened load durations and reduced the impurity burden per cycle, although productivity decreased. Additionally, the CIP step was extended to improve cleaning, and the batch loading flow rate was reduced to 270.5 cm/h to match the longer elution/CIP duration. The final time schedule for design B is shown in **Figure 1E**.

The first CaptureSMB run using design B was successfully conducted over four cycles. Peak shape differences still appeared after cycle 2 but were considerably smaller than in design A. During this run, a pressure spike occurred before cycle 3, leading to an extended hold that may have promoted fouling (**Figure S3**). A second CaptureSMB run using design B was carried out as a repetition (**Figure 3 A** and **B**). Although pressure trends (**Figure S1**) and peak shape changes (**Figure 3 B** and **C**) still indicated gradual accumulation of impurities, the yield remained stable over the cycles for both design B runs (**Figure 4A**). Compared to the batch runs, the recovery was on average +14.3% higher for CaptureSMB operation (**Figure 4B**).

**Figure 3:**
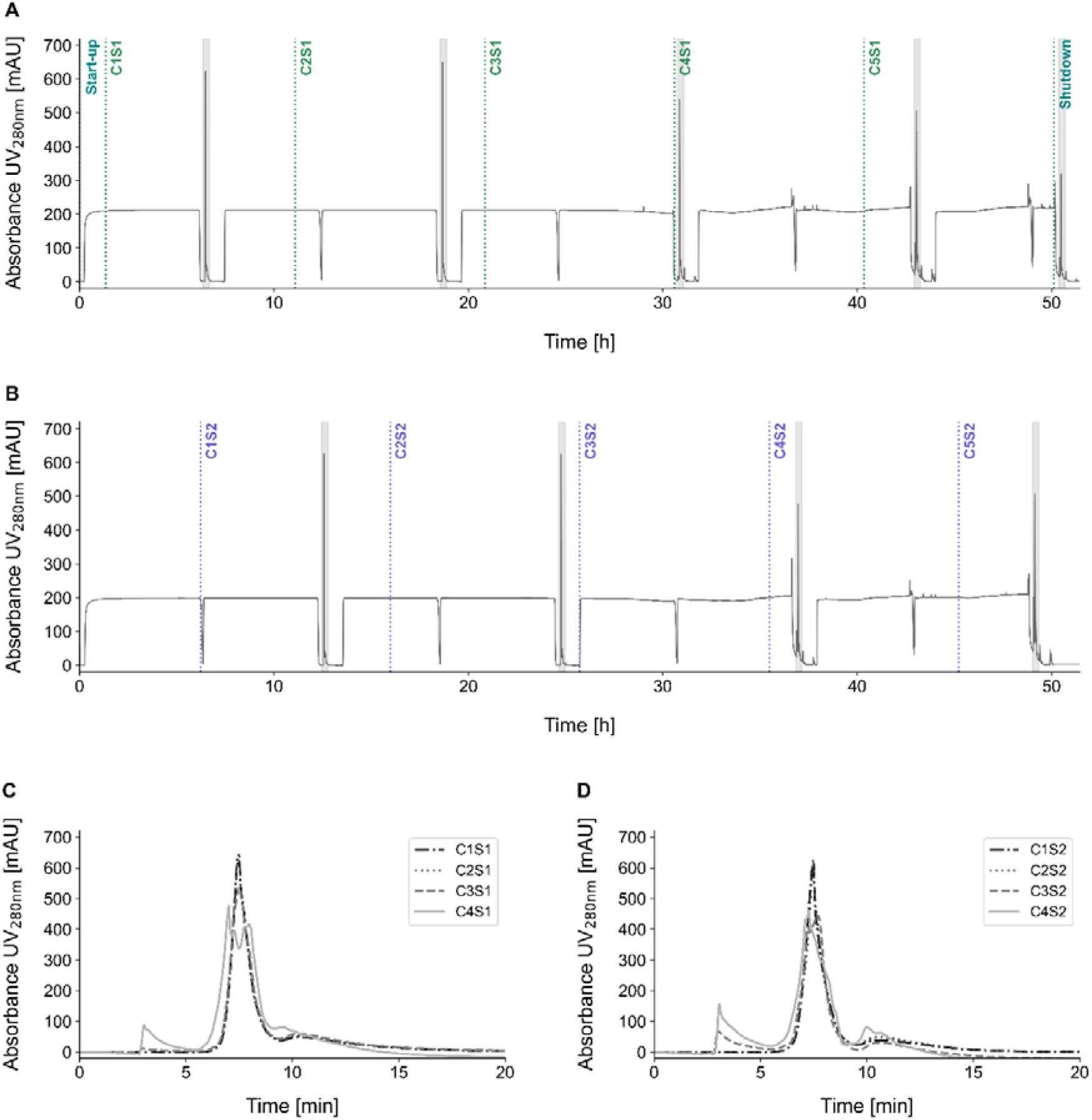
Chromatograms of the second CaptureSMB run following design B, for the capture of secreted rAAV5. The UV_280nm_ signals for column 1 **(A)** and column 2 **(B)** are shown, with the elution phases indicated in grey shading. The superimposed elution peaks of the four cycles are shown for column 1 **(C)** and column 2 **(D)**. C: cycle; S: switch.

**Figure 4:**
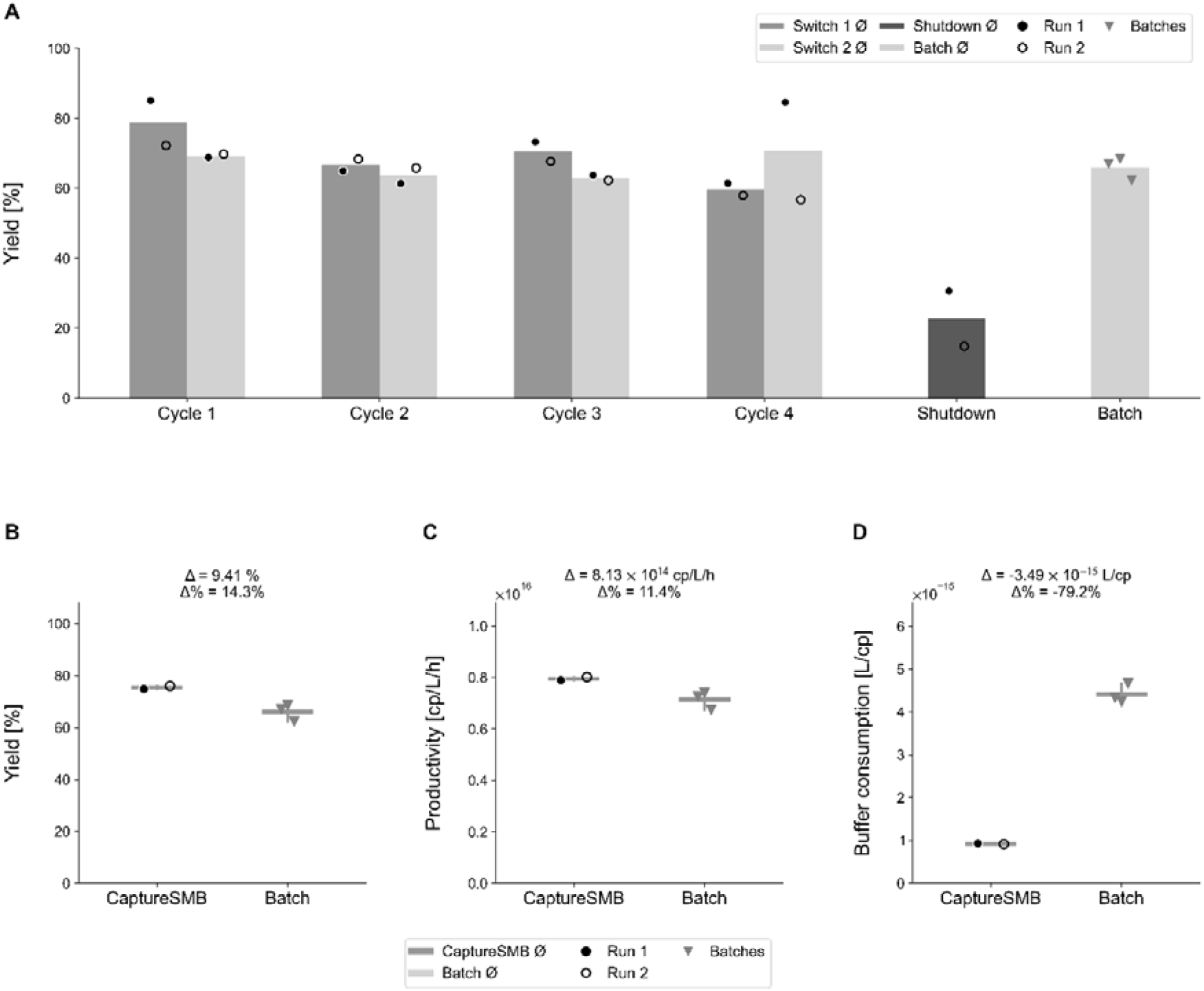
Process performance comparison of CaptureSMB and batch operations. **(A)** Yield over four cycles for two CaptureSMB runs following design B, including shutdown and batch references. Bars represent mean values. **(B)** Comparison of yield, **(C)** productivity, and **(D)** buffer consumption between CaptureSMB design B and batch processes. In all panels, filled circles (●) indicate individual measurements for CaptureSMB run 1, open circles (○) indicate CaptureSMB run 2, and triangles (▾) indicate individual batches. In (B - D), horizontal lines represent group means and vertical lines show the observed range. The reported Δ values correspond to the absolute difference in mean performance between CaptureSMB and batch processes (CaptureSMB - batch), while Δ % represents the relative difference normalized to the batch mean.

The process performance was further assessed by comparing the productivity and buffer consumption between the CaptureSMB and the three batch runs as described in the material and methods (**Equation 1**, **Equation 2**, **Equation 3**). Continuous operation improved productivity by +11.4% and reduced buffer consumption by -79.2% (**Figure 4 C** and **D**, **Table S1)**. Even with only four cycles and the need to reduce the interconnected loading, CaptureSMB was able to outperform batch operation.

As the altered peak shape after two cycles could indicate an accumulation of impurities in later cycles, critical quality attributes of the elution pools were analyzed individually in more detail. Overlaid analytical peaks of the AEX-HPLC analysis for UV 260 nm showed almost identical profiles for individual switches, indicating consistent recovery per elution (**Figure. S4**). The full capsid ratio remained constant across all CaptureSMB and batch eluates (23.2 to 24.4%, **Figure 5A**). Early CaptureSMB eluates showed approximately one order of magnitude lower DNA levels compared to the batch processes (**Figure 5B**). Thereafter, the residual DNA level increased in cycle 3 and 4 to levels comparable to the batch process. For the normalized total protein, the CaptureSMB eluates showed lower amounts than the batch runs (**Figure 5C**). In the continuous CaptureSMB process, the calculated protein content per viral genome (µg/vg) was lower than in the batch runs. This can be explained by the much larger amount of viral genomes recovered within the same elution volume in the continuous mode, while the amount of co-eluting protein impurities did not increase proportionally. As a result, the virus content is higher relative to the baseline of protein impurities, leading to lower µg of protein per vg. Nevertheless, an upward trend was still apparent, with the shutdown sample approaching the values observed for the batch runs. Since the load density during shutdown is comparable to that of a conventional batch operation, the higher normalized total protein content, reaching levels close to those of the batch process, is consistent with expectations.

**Figure 5:**
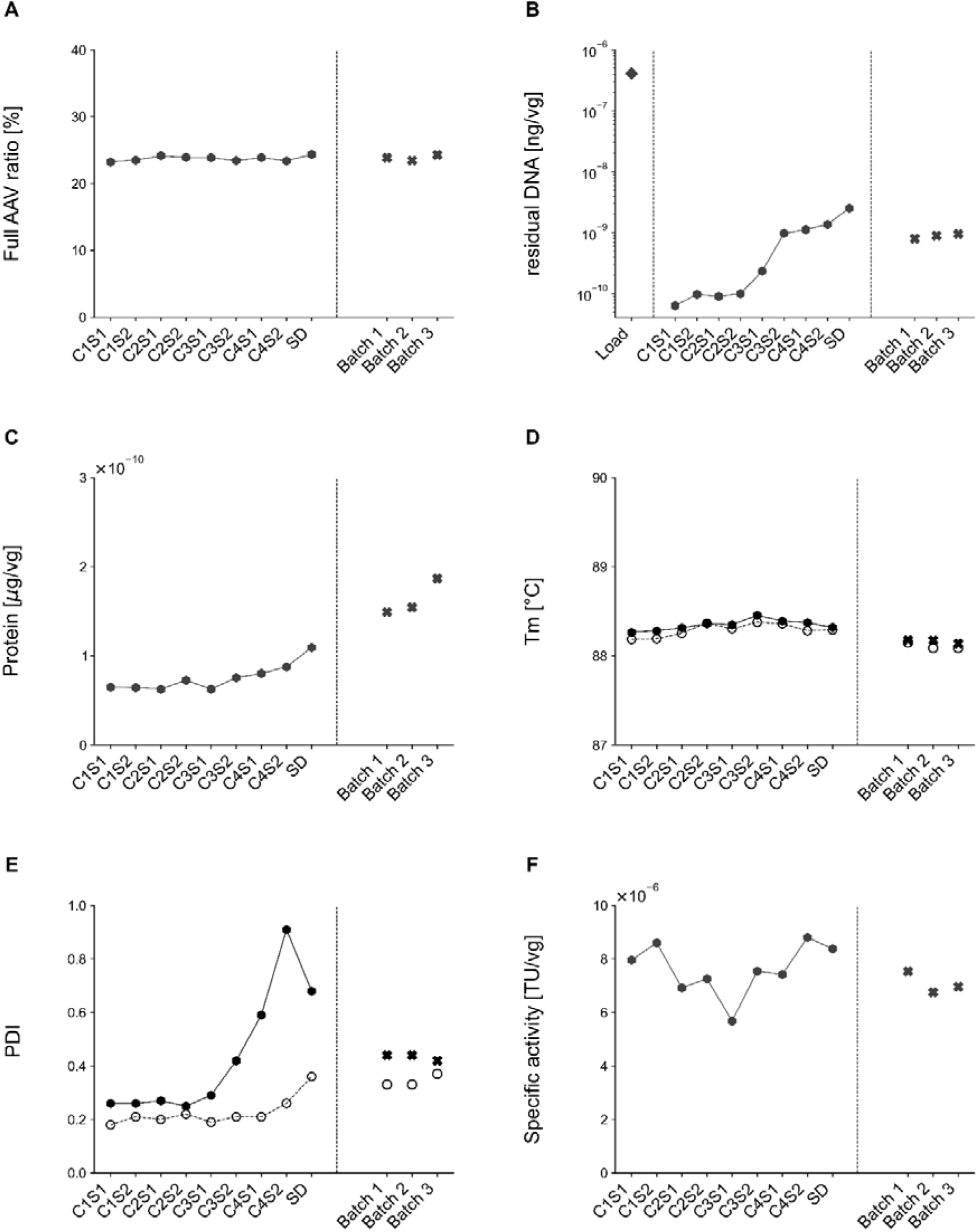
Comparison of the quality attribute between the CaptureSMB and batch chromatography. For the comparison, the eluates of the second CaptureSMB run following design B were analyzed and compared to the batch eluates (N = 3). The attributes measured were the **(A)** full% measured by AEX-HPLC, **(B)**, residual DNA per vg, **(C)** total protein per vg, **(D)** T_m_ and **(E)** PDI with the full dots pre- and the white dots post 0.2 µm filtration, and **(F)** the specific activity. C: cycle; S: switch; SD: shutdown.

Furthermore, the purified rAAV5 particles showed high thermal stability, with melting temperatures ranging from 88.1 to 88.5 °C (**Figure 5D**) which is consistent with previous reports identifying rAAV5 as the most thermally stable among common rAAV serotypes (Bennett et al., 2017; Lins-Austin et al., 2020; Pacouret et al., 2017; Rayaprolu et al., 2013).

The PDI values obtained by DLS showed a pattern similar to that observed for residual DNA (**Figure 5E**). From cycle 3 onwards, the PDI increased rapidly. In the early CaptureSMB cycles, PDI values were lower than those observed for the batch process, whereas in the later cycles they exceeded the batch values. After 0.2 µm filtration, the PDI generally decreased slightly, with a more pronounced reduction in the later cycles where the initial values were higher.

Finally, the transduction efficiency of the individual eluates was analyzed (**Figure 5F**). Throughout the entire CaptureSMB process, the specific activity was stable. Compared to the batch process, the CaptureSMB eluates showed a slightly higher average specific activity, although no statistical meaningful calculation could be performed given that several more experiments are required to calculate appropriate statistical tests. Thus, the continuous process maintained the activity of the recovered rAAV5.

## Discussion

The improvement in yield in CaptureSMB is relevant in the context of rAAV manufacturing, where losses in downstream steps substantially affect overall process performance. Increased resin utilization under continuous operation allows higher loading densities than conventional batch chromatography, where conservative underloading is often required to avoid premature product loss. Given the high cost of rAAV affinity resins, improved utilization directly translates to reduced COG, making higher loading densities particularly advantageous. Because these resins frequently show improved recovery at higher loading, CaptureSMB appears especially favorable and may contribute to the higher overall recovery observed.

This advantage is accompanied by gains in productivity and decreases in buffer consumption, even with only four cycles. Longer cyclic operations are expected to exhibit substantial higher gains, since start-up and shutdown phases are typically the least efficient steps in continuous chromatography. The start-up is required to equilibrate the column and pre-load the first column to the defined batch load density. At the end of the process, the shutdown serves to recover the remaining product from the last column loaded during the batch loading step. In terms of buffer consumption and product recovery, both phases are similar to a conventional batch process and therefore reduce the overall efficiency of the continuous setting. Thus, the observed advantage of continuous chromatography based on only four cycles suggests that further efficiency gains are possible with increased resin durability and more cycles.

The altered peak shape after two cycles, the gradual increase in column pressure drop, and increased residual DNA indicate fouling from nonspecific DNA binding not fully removed by CIP, as reported in earlier studies (Leibiger et al., 2024; Soni et al., 2025, 2024). The upward trend in DNA was also observed in batch operation, although to a lesser extent per repetition. A similar pattern was observed for PDI data. Elevated PDI in later cycles of CaptureSMB could indicate the formation of larger particles or aggregates, potentially involving rAAV–DNA complexes, as previously reported by Wright et al. (2005). Based on our observations it is plausible that such large complexes formed between rAAV particles and residual DNA or other contaminants may impede unhindered transport and binding within the column and contribute to fluctuations in the calculated breakthrough. This effect may be more pronounced in low-titer feed streams, where the impurity burden per unit of product loaded is higher, promoting the buildup of DNA–rAAV aggregates during breakthrough operation. The resulting interactions are likely driven by complex feed–resin dynamics that are not fully captured by simplified chromatographic models. While the underlying mechanisms were not resolved in this study, such behavior is plausible in crude harvest-derived rAAV purification, where residual DNA, proteins and other contaminants may influence transport and binding.

Accordingly, the robustness of cyclic operation highly depends on feed composition. Although protein impurities are removed efficiently, DNA remains the primary suspected fouling cause. Since no nuclease was used and no preceding TFF was performed prior to loading, the columns were exposed directly to untreated feed. Under these challenging conditions, cycle-dependent increases in residual DNA levels and higher PDI values in later cycles indicate that fouling or progressive accumulation of impurities occurred during repeated column use. These observations suggest that stable cyclic operation depends not only on the applied rAAV load, but also strongly on the impurity burden of the crude feed relative to the product concentration.

Higher-titer material, with lower impurity/product ratios, showed a lower level of fouling. Although these observations were not part of the scope of this study, they are consistent with the hypothesis that higher impurity/product ratios in harvest place a greater burden on the affinity matrix and can reduce process stability over repeated cycles.

From an industrial perspective, these findings are particularly relevant because affinity chromatography and early-stage clarification are associated with the highest downstream COG and longest operating times in rAAV workflows (Sarkis et al., 2023; Thakur et al., 2024). Under conventional batch operations, capture columns are usually substantially oversized simply to avoid overly long loading times when handling high-volume harvests (Lyle et al., 2024). This oversizing represents an economic bottleneck, as the high-cost affinity ligands are underutilized (Chu et al., 2023). In practice, current good manufacturing practice (cGMP) performance is further limited by feedstock impurities. Clarified lysates often contain high levels of host cell proteins and rAAV–DNA complexes, which cause rapid fouling of the column (Soni et al., 2024). Managing fouling requires balancing two competing needs: ensuring effective cleaning while avoiding excessive chemical damage to the ligand. Consequently, CIP protocols must be tailored carefully. Lower-intensity cleaning conditions require longer contact times to ensure sufficient cleaning depth without accelerating ligand degradation. However, regardless of optimization efforts, the combined effects of repeated chemical exposure and gradual impurity accumulation mean that validated multi-use lifetimes in batch mode are typically limited to fewer than 20 runs (Chu et al., 2023).

Shifting to a continuous multi-column configuration offers a clear operational advantage. This approach enables rapid, efficient loading across multiple smaller columns, removing the need for oversizing (Lyle et al., 2024). By maximizing the target breakthrough percentage under these intensified conditions, resin use is optimized, and buffer consumption is reduced before cleaning-related performance declines. In this way, manufacturers can overcome the traditional batch trade-off between speed and column size, thereby reducing the costs associated with short resin lifespans.

The productivity increase observed even with the relatively low number of cycles performed in this study demonstrates that CaptureSMB can deliver clear benefits in productivity and buffer consumption. The potential additional gains with a higher cycle count were estimated by evaluating the cyclic operation only, without start-up and shutdown phases. In this scenario, productivity increased by +18% and buffer consumption dropped by -82% (**Table S1**). Nevertheless, fouling in design A required process adjustments for the final design B, and these modifications also reduced overall process performance. Given that current rAAV affinity resins are still far less stable and cleanable than modern protein A resins (Chu et al., 2023), it is reasonable to expect that the benefits seen here could be amplified in future when resin stability and cleaning efficiency are improved. The extended loading times in continuous CaptureSMB operation did not compromise capsid integrity, as shown by the high thermal stability (T_m_ ∼88 °C) of purified rAAV5 particles. Most importantly, the transduction efficiency of the recovered rAAV5 was maintained throughout the continuous process, highlighting the robustness of this approach.

In addition, the simplified midstream strategy may offer broader process advantages across the rAAV workflow. By avoiding full cell lysis, the process may reduce the release of host cell-derived impurities into the harvest. It also eliminates the need for a TFF step before capture, thereby removing an entire unit operation. Likewise, omission of nuclease treatment may reduce process complexity and cost (Thakur et al., 2025a). Residual DNA contributed to fouling under low-titer conditions in our study. In experiments with higher AAV titers, where the impurity-to-product ratio was reduced, fouling was not observed. This indicates that the impact of residual DNA on fouling depends strongly on the relative impurity burden. Taken together, this approach allows a simplification of the overall rAAV production workflow.

## Conclusion

This work evaluated CaptureSMB as an alternative to conventional batch affinity chromatography for direct capture of rAAV5 from crude harvest. Compared with batch operation, CaptureSMB achieved higher productivity (+11.4%), increased product recovery (+14.3%), and strongly reduced buffer consumption (-79.2%). These advantages of CaptureSMB over conventional batch operation stem from the improved resin capacity utilization and elevated load flow rates with minimized product loss. Critical quality attributes including full/empty ratios, and transduction efficiency remained consistent throughout all cycles whereas DNA and protein contaminations remained below batch values.

Overall, the twin-column CaptureSMB process emerges as a promising strategy for intensifying the affinity capture of rAAV5 directly from untreated harvest, reducing the number of pre-treatment steps that typically reduce yield. Under the tested conditions, it enhances productivity and recovery while reducing buffer consumption relative to batch operation. While conceptionally ready for scale-up, successful implementation depends on feed quality and robust control of impurity-driven fouling during cyclic operation. By enabling direct capture with high performance and reduced resource consumption, this approach also lays the technological groundwork for future fully integrated continuous manufacturing of viral vectors.

## Author contributions

**Julia M. Müller**: Conceptualization, methodology, experimental work, data analysis, writing – original draft, review & editing. **Daniela Tobler**: Experimental work. **Jules Bühler**: Experimental work. **Damian Hauri:** Experimental work – review. **Richard Plieninger:** Experimental work. **Sven Göbel:** Experimental work, writing – review and editing. **Yuki Higuchi**: Conceptualization and writing – review and editing. **Ryosuke Takahashi:** Conceptualization and writing – review and editing. **Sebastian Vogg**: Conceptualization and writing – review and editing. **Thomas Müller-Späth**: Conceptualization, project administration, funding acquisition, and writing – review and editing. **Thomas K. Villiger**: Conceptualization, project administration, funding acquisition, and writing – review and editing.

## Acknowledgements

The authors would like to thank Patrick Werder for his excellent technical support and providing the rAAV plasmids. We are also grateful to Stefanie Reiter for her dedicated analytical support. Moreover, we would like to thank Levitronix and Asahi Kasei for providing equipment, such as the PuraLevi30 pumps and MF-SL hollow-fiber modules.

## Abbreviations

(r)AAV: (Recombinant) Adeno-associated virus
AEX: Anion exchange
ASSB: Assay buffer
BCA: Bicinchoninic Acid
bGH: Bovine growth hormone
BT: Breakthrough
cGMP: Current good manufacturing practice
CIP: Cleaning-in-place
CMV: Cytomegalovirus
COG: Cost of goods
Cp: Capsid particles
CV: Column volumes
DBC: Dynamic binding capacity
DLS: Dynamic light scattering
(ds)DNA: (double stranded) Deoxyribonucleic acid
DNase: Deoxyribonuclease
(E)GFP: (Enhanced) green fluorescent protein
ELISA: Enzyme-linked immunosorbent assay
Em: Emission
Ex: Excitation
HEK293F: Human embryonic kidney 293 cells
HPLC: Highperformance liquid chromatography
MOI: Multiplicity of infection
PDI: Polydispersity index
PEI: Max Polyethyleneimine Max
pHelper: Helper plasmid
PI: Plasmid insertion designation
PLW: Post-load wash
pRep/Cap: Replication/capsid plasmid
pTransfer: Transfer plasmid
SMB: Simulated moving bed
(r)TFF: (reverse) Tangential flow filtration
T_m_: Melting temperature
UV: Ultraviolet
VCC: Viable cell concentration
Vg: Vector genomes
WPRE: Woodchuck Hepatitis Virus Posttranscriptional Regulatory Element

