## Supplementary Information for "Continuous Capture of recombinant AAV Particles Using Twin-Column CaptureSMB"

Hofackerstrasse 30

4132 Muttenz

Switzerland

**Table S1:** Process performance parameters – Batch versus CaptureSMB. The two CaptureSMB runs following design B were evaluated and compared to three conventional batches. The productivity and buffer consumption were calculated once for the whole run, and once for the cyclic operation only.

| | | Total particles recovered | Yield | Productivity | | Buffer consumption | |
|---|---|---|---|---|---|---|---|
| | | | | Total run | Cycles | Total run | Cycles |
| | | [cp] | [%] | [cp/L/h] | | [L/cp] | |
| CaptureSMB | Desing B, run 1 | $3.98 \times 10^{14}$ | 74.8 | $7.89 \times 10^{15}$ | $8.32 \times 10^{15}$ | $9.25 \times 10^{-16}$ | $8.01 \times 10^{-16}$ |
| | Design B, run 2 | $4.04 \times 10^{14}$ | 76.0 | $8.01 \times 10^{15}$ | $8.45 \times 10^{15}$ | $9.11 \times 10^{-16}$ | $7.88 \times 10^{-16}$ |
| | Ø | $4.01 \times 10^{14}$ | 75.4 | $7.95 \times 10^{15}$ | $8.39 \times 10^{15}$ | $9.18 \times 10^{-16}$ | $7.94 \times 10^{-16}$ |
| Batch | Run 1 | $9.05 \times 10^{12}$ | 67.0 | $7.25 \times 10^{15}$ | | $4.33 \times 10^{-15}$ | |
| | Run 2 | $9.25 \times 10^{12}$ | 68.5 | $7.41 \times 10^{15}$ | | $4.24 \times 10^{-15}$ | |
| | Run 3 | $8.42 \times 10^{12}$ | 62.4 | $6.74 \times 10^{15}$ | | $4.66 \times 10^{-15}$ | |
| | Ø | $8.90 \times 10^{12}$ | 65.4 | $7.13 \times 10^{15}$ | | $4.41 \times 10^{-15}$ | |

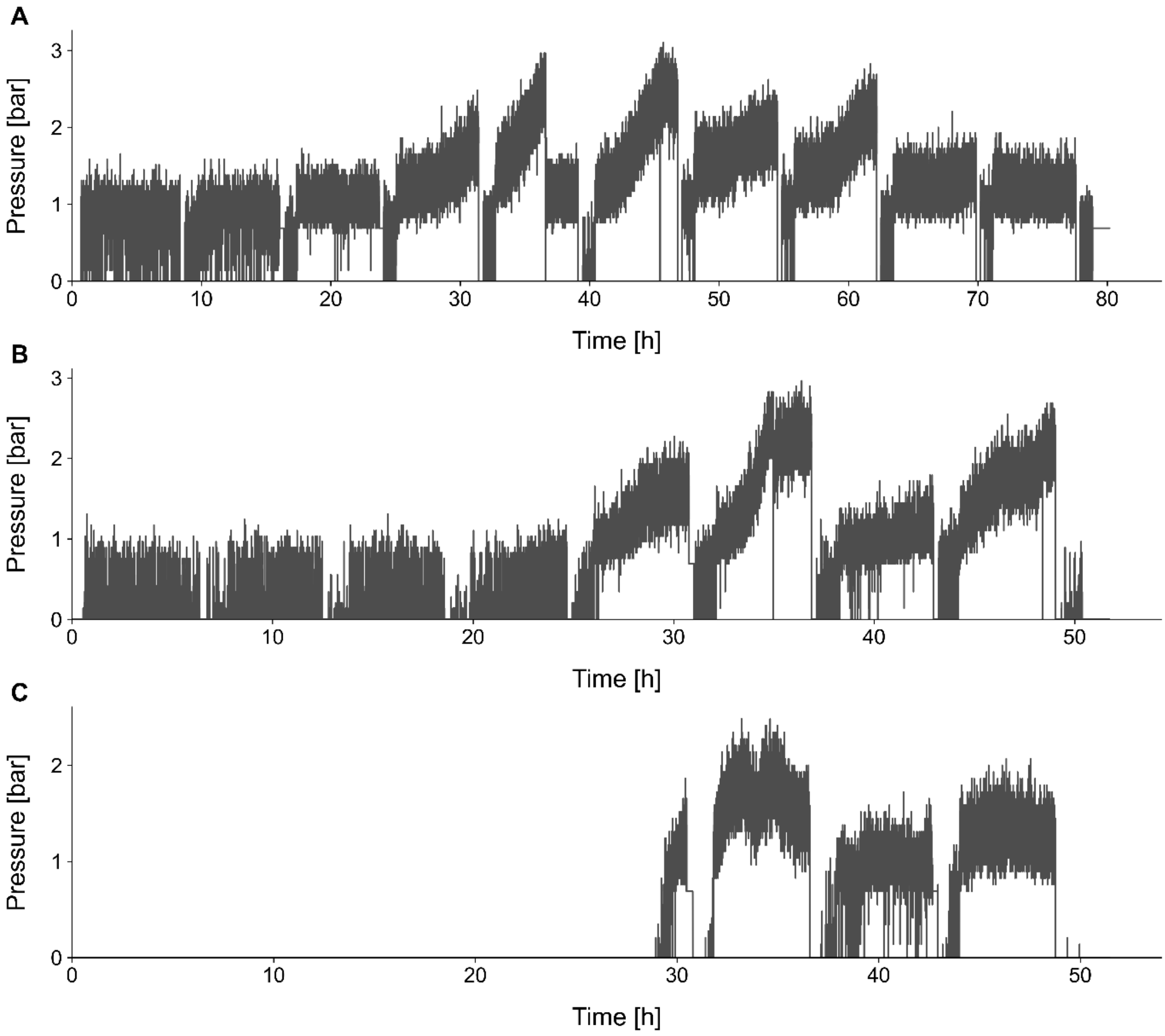


**Figure S1:** Continuous sample pump pressure profiles of the rAAV5 CaptureSMB runs. Concatenated pressure data for CaptureSMB run 1 of design A (**A**), and the run 1 (**B**) and run 2 (**C**) of design B.

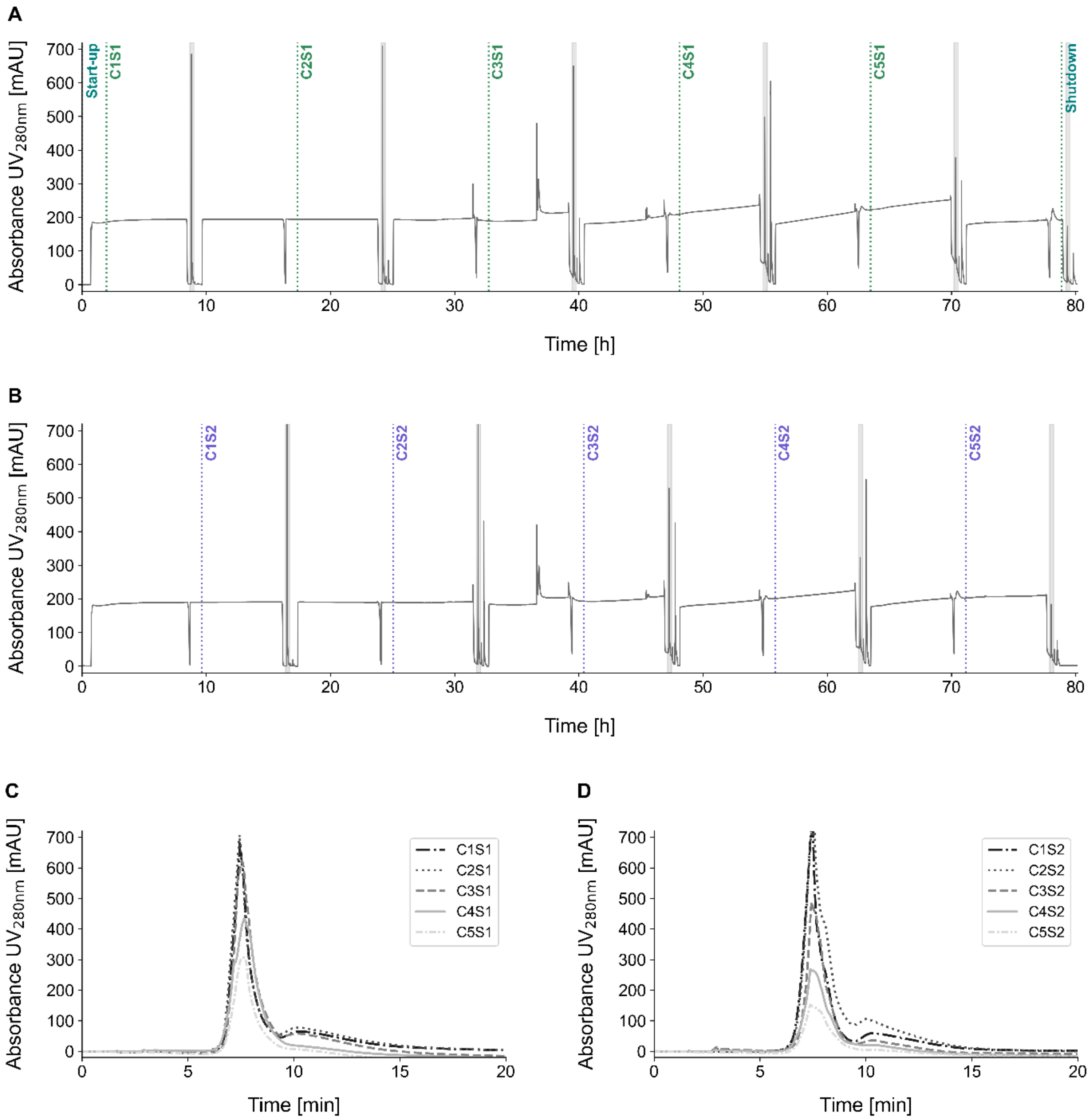


**Figure S2:** Chromatograms of the first CaptureSMB run following design A, for the capture of secreted rAAV5. The $UV_{280nm}$ signals for column 1 (**A**) and column 2 (**B)** are shown, with the elution phases indicated in grey shading. The superimposed elution peaks of the five cycles are shown for column 1 (**C**) and column 2 (**D**). C: cycle; S: switch.

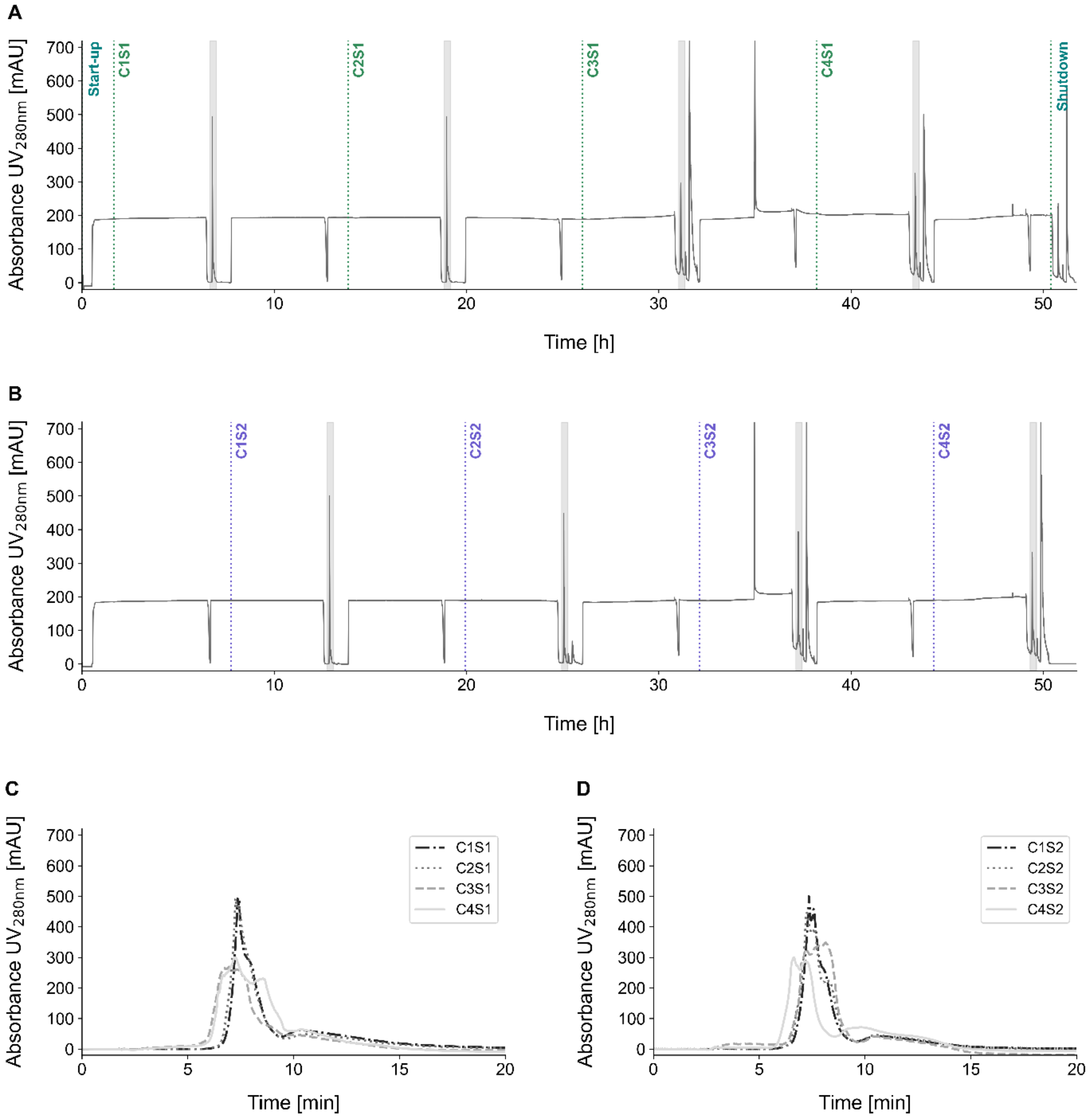


**Figure S3:** Chromatograms of the first CaptureSMB run following design B, for the capture of secreted rAAV5. The $UV_{280nm}$ signals for column 1 (**A**) and column 2 (**B)** are shown, with the elution phases indicated in grey shading. The superimposed elution peaks of the four cycles are shown for column 1 (**C**) and column 2 (**D**). C: cycle; S: switch.

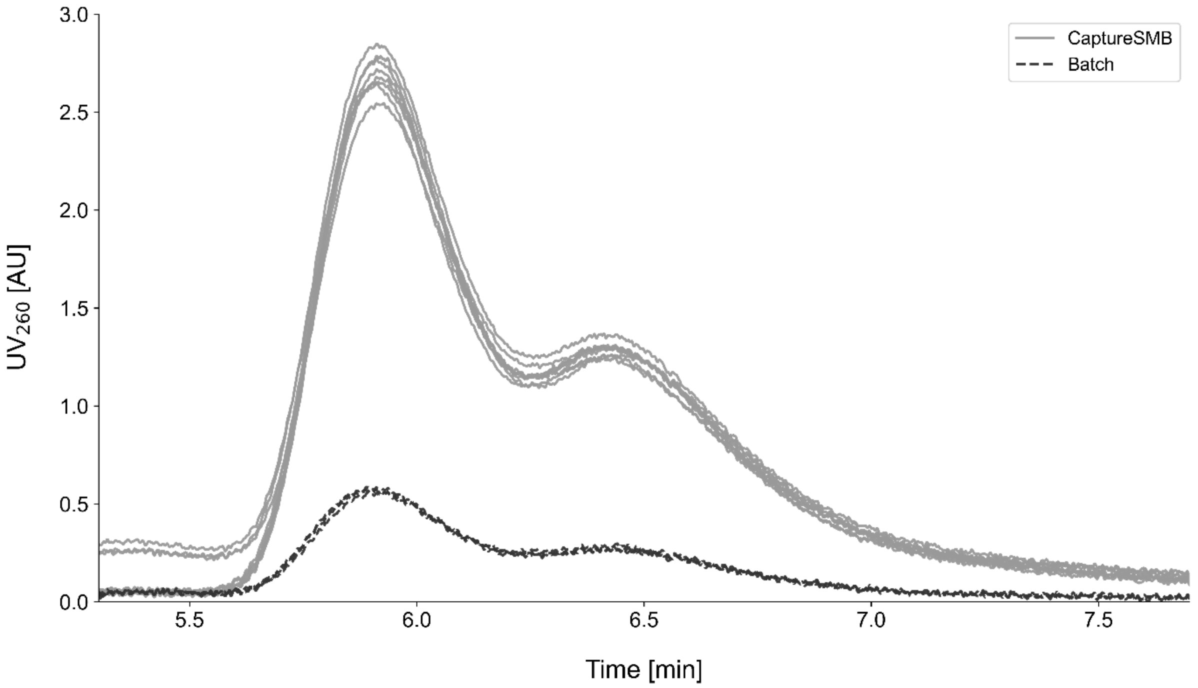


**Figure S4:** Chromatogram of the AEX-HPLC full rAAV5 % analysis for the individual eluates of the CapturеSMB run 2 of design B, in comparison to the batches. The peak at 5.9 min corresponds to the empty rAAV5 particles, and the peak at 6.4 min represents the full rAAV5 particles.
